# Low-cost and versatile electrodes for extracellular chronic recordings in rodents

**DOI:** 10.1101/2020.02.06.937201

**Authors:** Arthur S C Franca, Josephus A van Hulten, Michael X Cohen

## Abstract

Electrophysiological data are used to investigate fundamental properties of brain function, its relation to cognition, and its dysfunction in diseases. The development of reliable and open-source systems for electrophysiological data acquisition is decreasing the total cost of constructing and operating an electrophysiology laboratory, and facilitates low-cost methods to extract and analyze the data (Siegle et al., 2017). Here we detail our method of building custom-designed low-cost electrodes. These electrodes can be customized and manufactured by any researcher to address a broad set of research questions, further decreasing the final cost of an implanted animal. Finally, we present data showing such an electrode has a good signal quality to record LFP.

## 1.0 Introduction

The extraction of extracellular voltage fluctuations, from spiking activity to neuronal oscillations, is one of the most widely used techniques in neuroscience (Siegle et al., 2017). Electrophysiological recordings are crucial to relating dynamics across different spatial scales - from individual spiking to the activity of thousands of neurons in the electroencephalogram - and across different temporal scales - from milliseconds to hours. Such type of data has been instrumental in insights and discoveries about cognition and brain functions (Cohen, 2017).

A modern electrophysiology recording system must allow for multichannel recordings with adaptable geometry for recording from multiple brain regions of varying depth and curvature, and must be easily integrated into an optogenetics or chemogenetics delivery system. Two classical ways to approach this are to use “hyperdrives” - carrier structures that enable the movement of individual electrodes or bundles of electrodes such as tetrodes (Battaglia et al., 2009; Bragin et al., 2000; Brunetti et al., 2014; Kloosterman et al., 2009; Liang et al., 2017; McNaughton, 1999; Michon et al., 2016), or to use silicon probes (Bragin et al., 2000; Jun et al., 2017; Lopes-dos-Santos et al., 2018; Ulyanova et al., 2019). Both approaches have several advantages including high-density recordings or layer-specific profiles. However, both of these existing methods also have disadvantages, including the size of the structure in and on the animal’s head, limited number of simultaneously accessible brain regions, a significant amount of time to build the apparatus, or the high price of commercially available electrodes.

We aim to describe a low-cost and versatile electrode system for extracellular chronic recordings in rodents. The electrodes described here are optimal for recording the Local Field Potential (LFP) and multi-unit activity across multiple brain regions simultaneously. Our design is small and lightweight and does not impair movement, animal welfare, or behavioral tasks. In this paper, we describe systematically how to manufacture each component, including video and options for customization of the electrodes to target any specific research question. The approximate cost per 32-channel electrode implant is 50 euros and takes around three hours to build. A chronic implant will continue to provide high-quality LFP data for months.

## 2.0 Method

### 2.1 Animals

We have developed and implanted our electrodes in mice and rats, and we continue to record high-quality LFP data up to 4 months after implant. The data shown in this paper are from Black57 mice, recorded using implants in different brain regions for verification of the quality of signal to noise. All animals were recorded in their home cages, having free access to food and water. All experiments were approved by the Centrale Commissie Dierproeven (CCD) and it is according to all indications of animal welfare body [Approval number 2016-0079].

### 2.2 Material List

#### 2.2.0 Wires

Several types of materials can be used to manufacture electrodes. Here we chose to use Tungsten (99.5%) for the electrode due to the high resistance with very thin diameters, and stainless steel for the ground wire for both resistance and low cost. We use and recommend the usage of 50μm and 35μm tungsten wire (rods, not rolls). Wires smaller than 35μm increase the risk of bending during the implantation surgery. We purchased wires in bulk from <California Fine Wire Company>, although any other supplier would be suitable.

#### 2.1.2 Grid

The individual wires need to be aligned, and for this we use an alignment grid. The idea is to place two such grids with holes drilled at the same locations, one 3cm on top of the other. The individual wires are then fed through these holes. We manufacture these grids using a micro-manipulator and micro drills (~90μm tip) for drilling holes in plastic paper. We recommend a spacing of at least 250μm between the holes of the grid. If the spacing is lower than 200μm, the electrodes may form a compact surface, increasing the risk of damage to the brain during implantation surgery. A combination of two grids with the same specifications is used to align the electrodes (see Figure 1, <u>Video 1</u>).

**Figure 1:**
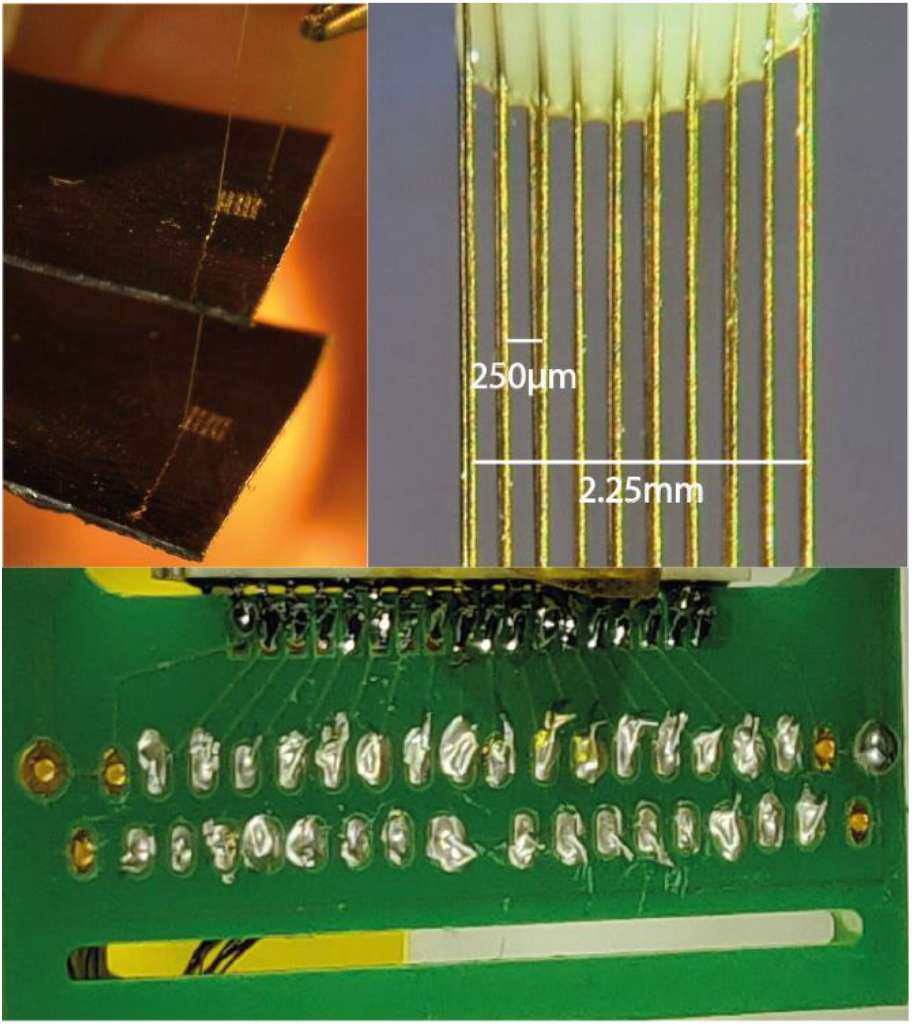
Different electrode arrays components: Top panel custom grid to align the electrodes 250μm apart. Bottom panel custom PCB during finalization phase, notice the silver paint to connect the wire to the PCB.

#### 2.1.3 Connectors

The manufactured electrodes presented here are based in the Intan/Open-ephys recording systems. The headstages used by these companies can be coupled with the Omnetics connector. It is therefore necessary to investigate which connectors are compatible with the headstage to be used. We use the connector A79026-001 (Omnetics Connector Corporation-USA). These connectors are compatible with the headstage RHD2132 for 32 channels.

#### 2.1.4 Printed Circuit Board (PCB)

For fixed implants, we use PCBs to link the electrodes to the connectors. The PCBs were designed using the free version of the software Eagle (https://www.autodesk.com/products/eagle/free-download). During the design of the PCB, the measure between the claws of connectors and also the space between the two lines of the connector have to be considered to avoid displacement of the Omnetic claws with the soldering points of PCB (<u>Video 1</u>). Details of the connectors and its measurements can be found on the Omnetics website. The PCBs were printed by Eurocircuits (https://www.eurocircuits.com/). Any company specialized in PCBs should be suitable.

### 2.2 Single wire arrays manufacturing

The process described below was video-recorded to facilitate visualization (<u>Video 1</u>). The video is currently available at mikexcohen.com/lowcostephys, and will be provided as a supplemental data file upon acceptance.

For single wire arrays, we used the components 2.1.1 to 2.1.4 described above. The fixed electrodes array can be implanted with or without the usage of PCBs. If Tungsten wires are used, we recommend PCBs due to the technical difficulties of soldering Tungsten directly to the connector. Therefore, we divide the manufacturing process into 3 steps:

1. *Electrode Alignment:* The electrode alignment is completely customized to any experiment to be planned. We here describe an implant optimized for multisite recordings in the medial prefrontal cortex. We designed the electrode to contain three rows of electrodes that would be implanted into one hemisphere. Using the grid of 250μm we designed the array to have a 3×10 geometry. The electrodes are laid into the grid in each row of 3, repeating 5 or 10 times, depending on whether a 32 or 16 channel connector is used. The tips of the electrodes can be aligned using a calypter (<u>Video 1</u>). It is important to consider the dorsal-ventral coordinate of the target site (in our example: 1.5mm), and add 2 or 3mm to prevent the PCB from touching the skull during the implant (<u>Video 1</u>).
2. *Soldering*: The PCB and Omnetics connectors should be placed in an aligned position considering the claws of the connector and the Soldermask of the PCB (<u>Video 1</u>). Due to the very small components to be soldered, we use a soldering tip of 200μm. The ground wire can be soldered directly to the PCB, using an appropriate flux for soldering stainless steel. Note that the ground wire can be directly soldered to a screw, or left to be connected to the screw during surgery.
3. *Assembling:* To finalize the electrode, the two processes mentioned above must be assembled. The aligned electrodes were glued to the PCB/connector. We recommend using photo-activated glue to accelerate the process. After that, each electrode is placed in the individual holes of PCB. At this point, it is important to keep track of the placement of each electrode for proper mapping of data channel to physical location. The excess wire is cut and the wire coat is removed with a surgical blade. Silver paint is applied to connect the tungsten wire to the PCB (Figure 1). The electrode is finalized using Epoxy glue to cover and protect the arrays and the connection between the electrodes and PCB is made by the silver paint (<u>Video 1</u>).

After finishing the procedure with 32 electrodes, the construction weighs 1.0 ~ 1.2g, corresponding on average less than 4% of the body weight of the mice or 0.5% of the rat body weight. The weight will be less for 16 electrodes.

### 2.3 Surgical procedure

The surgical procedure varies according to the type of electrode and target brain regions. We here describe a general procedure that should be adapted to the specific needs of the project.

1 - Pre-implantation: In the pre-implantation phase we used the standard procedure of stereotaxic surgery. 10-16 week old mice were anesthetized with Isoflurane (induction at 5% Isoflurane in 0.5L/min O2; maintenance at 1-2% Isoflurane in 0.5L/min O2) [Teva]. For surgery, mice were fixed in the stereotaxic instrument [Neurostar Stereotaxic]. After shaving, the skin was disinfected with ethanol (70%). The local anesthetic Xylocaine (2%, Adrenaline 1:200.000 [AstraZeneca]) was injected subcutaneously at the incision site before exposing the skull. Peroxide (10~20% H2O2; [Sigma]) was applied to the skull with a cotton swab for cleaning and visualization of bregma and lambda. Holes for support screws are drilled in the edges of the skull (note that the placement of screws is dependent on the position of electrodes).

2 - The window(s) in the skull through which the electrodes are lowered into the brain are drilled specifically to accommodate the type of arrays to be implanted. These windows should be as small as possible, because more bone removed leads to less remaining skull surface area for anchoring the electrodes (Figure 2A). For example, two electrode arrays should be done with two small windows rather than one larger window (Figure 2A). To avoid contact between the dental cement and the brain, vaseline can be applied to the windows after the implant (Figure 2B). Electrodes and screws are fixated onto the skull with dental cement (Super-Bond C&B) (Figure 2C). Approximately 40 minutes prior to the end of the surgery, saline and analgesic (Carprofen injected subcutaneous) injected to facilitate recovery of the animal.

**Figure 2:**
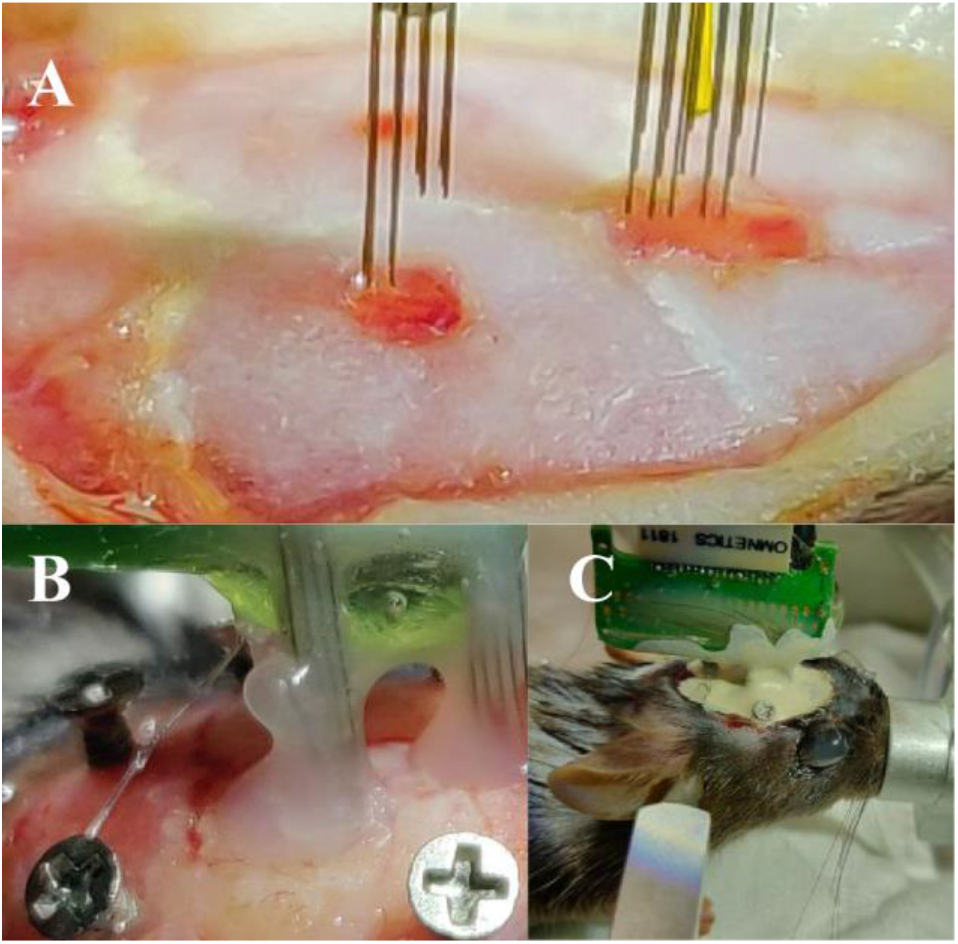
Electrode arrays implant surgery. Panels A, B and C shows different steps of the surgical implant. (**A**) Two craniotomies to accommodate arrays in different areas of the brain. (**B**) Vaseline to prevent the contact of dental cement with the brain. **(C)** Finalizing the surgery building the dental cement cap.

### 2.4 Data acquisition

Electrophysiology data were acquired using Open Ephys with a sampling rate of 30 kHz. Animals were recorded in their home cage for 10 minutes. During preprocessing, the data were downsampled to 1000 Hz, and EEGlab (Delorme et al., 2011) was used for visual inspection and cleaning artifacts. One mouse was kept implanted for 3 months to evaluate any changes in the signal characteristics.

### 2.5 Data Analysis

The data analysis was performed using Matlab (R2015b). Power spectrum density and the spectrogram representation was computed by the pwelch.m and spectrogram.m routines from the Signal Processing Toolbox. (Parameters: 50% overlapping, Hamming window of 6 s and Number of DFT points [2^13]).

### 2.6 Recycling materials

There is one component of the present electrode arrays that can be recycled: the connector, which is also the most expensive component of the array. After perfusion and brain extraction, the animal cap can be kept in acetone for 2 to 4 days. The acetone will dissolve the dental cement and epoxy used during the manufacturing. Finally, the soldering iron can be used to remove the connections between the connector and the PCB. In our experience, the connector can be reused twice without loss in the quality of the signal.

### 2.7 Electrode final cost

The price variation of each component used in the present manuscript can change according to the market value, the quantity of pieces purchased at a time (e.g., in bulk or individually), and the recycling capacity of the components. Therefore, the exact cost per electrode can only be approximated.

We order in bulk to produce around 60 electrodes arrays. The final cost for our array of 32 electrodes was 90 Euros, and the cost for a 16 channel array was 75 Euros. However, the most expensive component of the electrode - the connector - can be recycled. This factor brings the final cost to approximately 50 Euros/32Ch and 35 Euros/16Ch.

### 2.8 Manufacturing time

An inexperienced researcher or student might take 5 - 6 hours to build one electrode array. However, the learning curve is steep, and after some practice, it should take around 2-3 hours to construct a 32-channel array.

## 3.0 Results

To evaluate the quality of the signal that can be extracted with this type of electrode arrays, we recorded spontaneous LFP activity from an animal in their homecage in two different moments, day1 (First Day of recording after post-surgery recovery time) and 90 days after the first recording session to show that the implant sustains high signal quality for long periods of chronic recordings. We recorded from 3 independent regions of the brain to show the flexibility of the arrays.

Figure 3 presents raw signal traces (blue line) of three example channel from three different regions recorded in the same animal: hippocampus, mid-frontal cortex and posterior parietal cortex. Notice that the amplitude of the signal recorded in these channels is maintained after 90 days of recording. In the bottom panel, as expected, theta oscillatory activity can be verified in the hippocampus in the raw signal, which is translated in the power spectral density (PSD) and time-frequency spectrogram along the entire session of recording. Similar features can be verified after 90 days of recordings. Notice that after 90 days, different features can be observed in the LFP power; for example, by recording the animal in a new cage, novelty detection-related power in the beta2 band is apparent, as previously reported (Berke et al., 2008; França et al., 2014).

**Figure 3:**
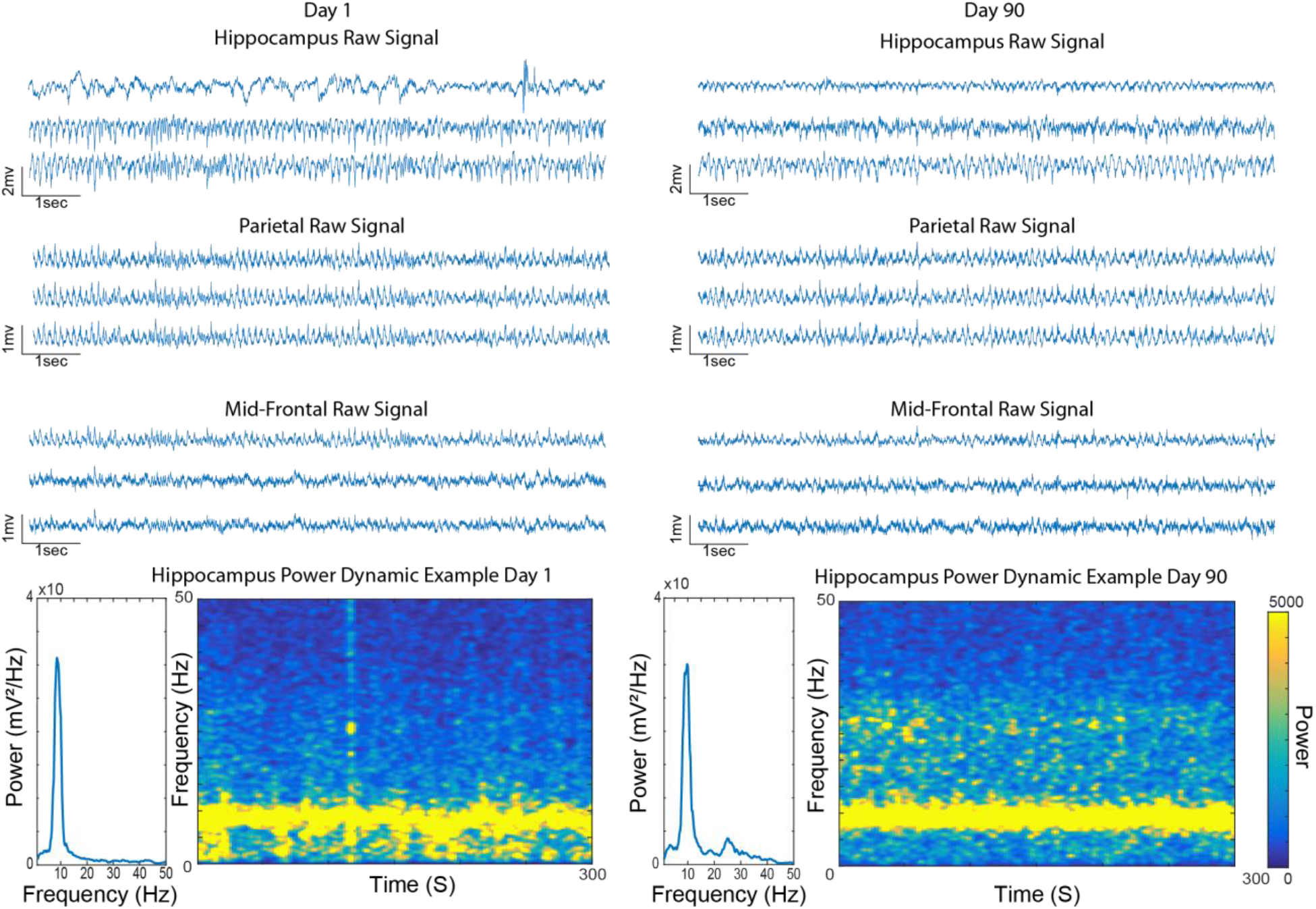
Signal quality maintained after 90 days of implantation. The figure shows examples of raw signal of the three regions recorded in the brain. Notice that the amplitude of the signal is maintained during 90 days of implantation. The bottom panel shows similar power dynamics characteristics in the theta band of frequency in both recording, day 1 and 90.

## 4.0 Discussion

The electrodes described here allow for addressing a wide range of research questions involving extracellular recordings electrophysiological data, with maximal flexibility at minimal cost. As demonstrated in the results section, we were able to extract LFP from several different regions simultaneously. We use these electrodes in freely moving animals, but they can also be used in acute experiments. Lastly, the arrays maintain integrity for long periods of time, allowing for investigations into long-term behaviors, learning, ageing, development, disease progression, and so on.

The compact shape of these electrodes, as well as the shape of the final implant in the animal (figure 2), are appropriate for recording during several types of behavioral experiments. Due to the light weight of the electrodes, even with multiple electrodes implanted in different brain regions, these electrode arrays do not interfere with animal welfare or behavioral performance (França et al., 2014). Besides that, these electrodes were used for long recording sections (up to 12 hours) without disturbing the sleep wake cycle of the animal (dos Santos Lima et al., 2019; França et al., 2015).

Although the electrode arrays are versatile to a range of applications, a limitation is that these arrays are not optimized for isolating spiking activity from individual units. This is partly due to the size of the electrode tip, and partly due to the fact that the electrodes are implanted without movable drives; thus, gliosis may prevent well-isolated spikes. This can be overcome by including micro-drives to move the electrodes, but this also increases the complexity of the implantation and also the total weight and size of the electrodes.

In conclusion, we have outlined a procedure to make very low-cost multielectrode arrays that are optimal for research questions that are based in LFP recordings and multi-unit cell activity, have the advantage of easily record from multiple regions of the brain, and are light enough to be used during behavioral performance. The electrodes are easy to build and easy to integrate into other open-source hardware and tools such as Open-Ephys.

